# Blocking plasmodesmata in specific phloem cell types reduces axillary bud growth in *Arabidopsis thaliana*

**DOI:** 10.1101/2020.12.28.424434

**Authors:** Andrea Paterlini, Delfi Dorussen, Franziska Fichtner, Martin van Rongen, Ruth Delacruz, Ana Vojnović, Yrjö Helariutta, Ottoline Leyser

## Abstract

The plasticity of above ground plant architecture depends on the regulated re-activation and growth of axillary meristems laid down in the axils of leaves along the stem, which often arrest as dormant buds. Plasmodesmata connecting plant cells might control the movement of regulators involved in this developmental switch. Constructs capable of occluding these structures were employed in phloem cell types, because of the importance of phloem in local and systemic trafficking. We show that over-accumulation of callose within companion cells of the Arabidopsis inflorescence reduces the growth rates of activated buds, but does not affect bud activation. Growth rate reductions were not dependent on the phloem-mobile strigolactone receptor, which regulates bud activation. Furthermore, there was no correlation with early bud sugar profiles, which can also affect bud activity and depend on phloem-mediated delivery. It is therefore possible that an as yet unknown mobile signal is involved in modulating branch growth rate.

## Main text

Shoot branching involves the coordinated regulation of the activity of meristems established in the axils of leaves along the stem (Mcsteen and Leyser, 2005). Once established, such axillary meristems often arrest as a dormant bud after the production of a few leaves. The hormone auxin, produced in the shoot apex, plays a central role in this process by moving downward in the stem and maintaining these axillary meristems in an inactive state, a process termed apical dominance (Morris, 1977, Snow, 1925, 1929). Since auxin does not enter the buds, the auxin transport canalisation model for bud regulation was postulated (Li and Bangerth, 1999; Bennett et al. 2006). According to this model, each bud, acting as an auxin source, must establish canalised auxin export in order to grow. The hormone self-reinforces its transport through positive feedback between flux and auxin transporter accumulation on the membrane of cells in the direction of flux (Sachs, 1969; Mitchison, 1980, 1981). Auxin largely relies on a series of active transporters, including of the PIN-FORMED (PIN) family for its cell-cell movement (Gälweiler et al., 1998; Bennett et al., 2014). Depending on the relative strengths of auxin sink in the stem and sources in the buds, and the level of feedback between auxin flux and transporter accumulation, some axillary buds might be able to activate while others would not (Prusinkiewicz et al., 2009).

In parallel to the systemic action of auxin, the transcription factors BRANCHED 1 (BRC1) and, to a lesser degree, its close paralogue BRC2, regulate shoot branching by operating at a local level within buds. They negatively regulate bud activation (Aguilar-Martínez et al., 2007) and act as signal integrators to adjust branching under a range of environmental conditions (Seale et al., 2017; Finlayson et al., 2010; González-Grandío et al., 2013). Further regulators of shoot branching, such as strigolactone and cytokinin hormones appear to act by modulating auxin transport canalisation and *BRC1* expression. These hormones, unlike auxin, enter the bud directly from the stem, moving in the xylem transpiration stream (Domagalska and Leyser, 2011).

Besides transmembrane transporters and flow in the lignified cells of the xylem, other regulators of bud growth could move via plasmodesmata (PD), the small channels connecting the cytoplasm of neighbouring plant cells (Li et al., 2020). Long distance transport could also occur in the phloem, a specialised conduit for nutrients and signals (Turgeon and Wolf, 2009). Two phloem mobile sugars, sucrose and trehalose 6-phosphate (Tre6P), have, for instance, been implicated in the control of shoot branching as their levels increase in buds upon apex decapitation (Fichtner et al., 2017; Mason et al., 2014) and Arabidopsis plants with altered levels of Tre6P show distinct branching phenotypes (Fichtner et al., 2020). Defoliation, removing the source of these compounds, or their exogenous application have opposing effects on bud growth (Fichtner et al., 2017; Mason et al., 2014). The role of these compounds is most likely a signalling rather than metabolic one as non-readily assimilable sugars still elicit growth effects (Barbier et al., 2015) and Tre6P is a known sucrose-specific signal in plants (Figueroa and Lunn, 2016). These metabolites can also influence PIN protein levels and *BRC1* expression (Barbier et al., 2015; Mason et al., 2014). The strigolactone receptor DWARF 14 (D14) is another macromolecule present in phloem sap of plants (Aki et al., 2008; Batailler et al., 2012) and its transport is necessary for tilling control in rice (Kameoka et al., 2016).

Genetic tools to modulate long distance and local cell-cell connectivity are available in plants, one of the more widely used being the *icals3m* system. This tool, of which we make use in this study, consists of a mutant version of a *CALLOSE SYNTHASE 3 (CALS3)* gene, engineered under the control of an estrogen transactivator and tissue specific promoters. Callose is a polysaccharide lining PD and its accumulation due to the mutant, over-active, enzyme results in occlusion of PD in a temporally and spatially controlled manner (Vaten et al., 2011).

The process of phloem unloading, the ultimate release of substances from this specialised conduit, has not been characterised in Arabidopsis buds. It could in principle occur symplastically (via PD) or apoplastically (via transporters) (Oparka et al., 1990). We imaged the *SUC2:GFP* reporter, driving free GREEN FLUORESCENT PROTEIN from the *SUCROSE TRANSPORT PROTEIN (SUC) 2* promoter, which is widely used to study symplastic unloading (Stadler et al., 2005; Imlau et al., 1999;) and observed a broad signal in sections across the inflorescence stem and its buds (Fig. 1a). The signal was beyond the domain of *SUC2* expression (Supp. Fig. 1a) so the result is compatible with a symplastic scenario. The fluorescence pattern is also similar to that observed in Arabidopsis roots at the tip, where PD-driven unloading is observed as a diffuse GFP signal, while higher up in the root the signal is restricted to the vasculature (in companion cells and transporting sieve elements) (Supp. Fig. 1b,c,d). To gain a quantitative appreciation of systemic delivery to the inflorescence we applied two tracers, 14C sucrose (Slewinski et al., 2009) and the fluorescent phloem mobile probe carboxytetraethylrhodamine (Knoblauch et al., 2015) to rosette leaves of plants grown axenically just after floral transition. The inflorescence was dissected into its component parts 16h later (Supplementary methods). Both tracers produced clear signals above background in all the organs of the inflorescence (Supp. Fig. 1e). The relative signals in the various organs (scaled by fresh weight and represented as percentage of total scaled signal) were largely equivalent between the two tracers with an exception in cauline leaves where more fluorescent probe was observed that 14C sucrose (Fig. 1b; Supp. Fig. 1g). This might be due to phloem to xylem transfer of the probe with cauline leaves, which experience the highest transpiration rate among the tissues sampled, accumulating most signal. Overall, the inflorescence stem and the shoot apex seemed to be stronger sinks than the buds, at least when the latter are small and dormant. Delivery of carboxytetraethylrhodamine, which is unlikely to be taken up by endogenous transporters, provides further evidence that unloading in the inflorescence is, at least in part, symplastic. The PD of vascular tissues would therefore play key roles in the process.

**Fig. 1.**
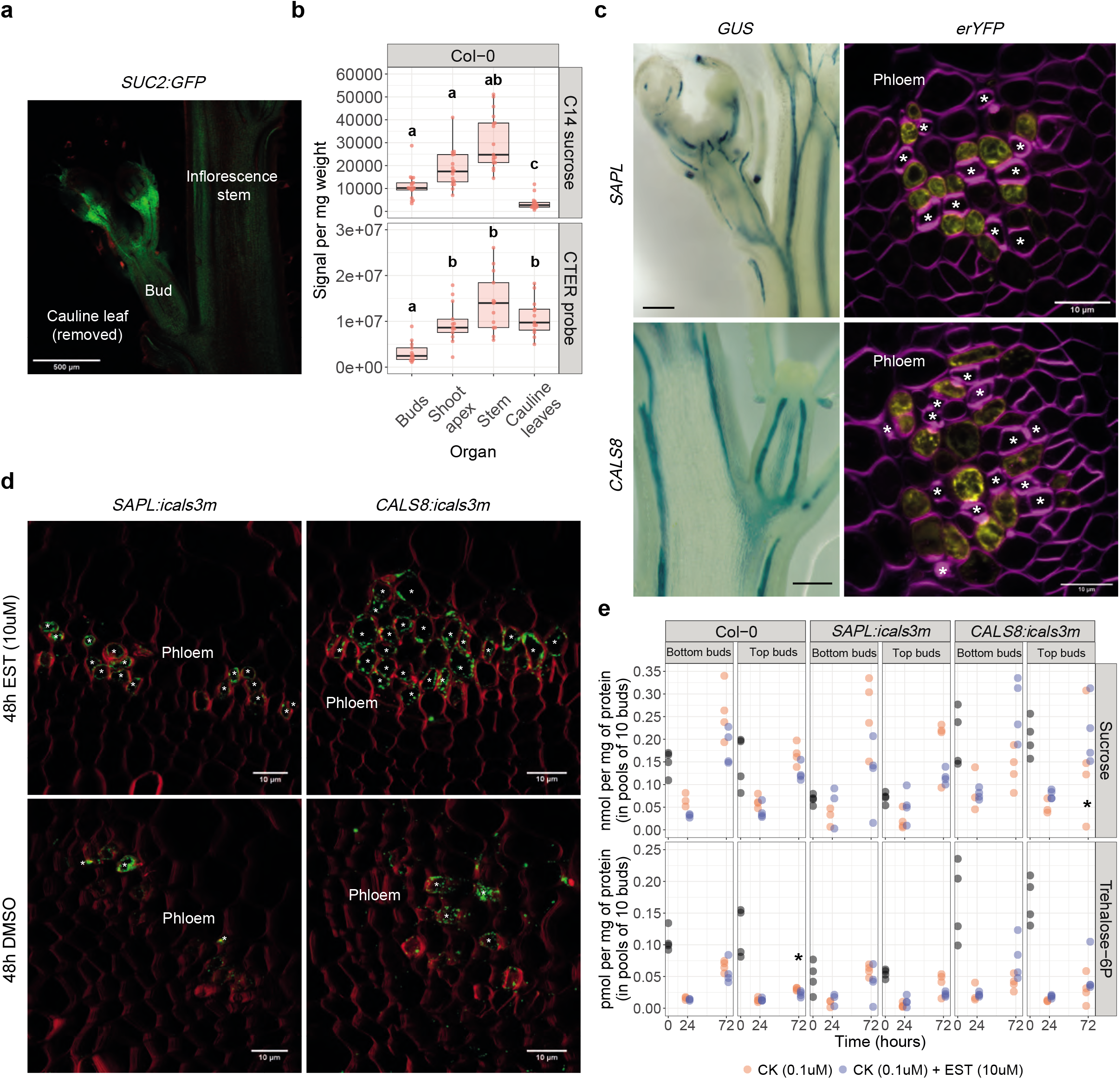
Phloem unloading and phloem cell-cell connectivity in inflorescence stems and buds. a. *SUC2:GFP* signal in a longitudinal section of the inflorescence stem and one of its buds. GFP is rendered in green, propidium iodine stain is false coloured in red. b. Radioactive (N=17) or fluorescence (N=13) signal intensity in inflorescence organs of plants supplied with label through the rosette leaves, scaled by fresh weight. Different letters indicate statistical differences in a Dunn’s test with a p-value threshold of <0.05. c. *SAPL/CALS8:GUS* signal in the inflorescence and *SAPL/CALS8:erYFP* fluorescent signal in the phloem part of vascular bundles of the inflorescence stem. YFP is rendered in yellow while calcofluor white stain is false coloured in magenta. Asterisks indicate sieve element (SE) cells. d. Callose immuno-labeling in inflorescence stem sections from explants supplied with DMSO or estradiol (EST) for 48h. Signal from secondary antibody is rendered in green while calcofluor white stain is false coloured in red. * indicates a cell with strong callose-related signal within the phloem. e. Sucrose and Trehalose-6P amounts at various time points after EST or non inductive treatment in buds of 2-node explants with intact apices. * indicates statistically significant differences in a two-tailed Mann-Whitney test with a p-value threshold of <0.05. N= 3-4 per timepoint, genotype and treatment. Scale bars are of 500um (a.), 100um (c. GUS images) and 10um (c. YFP images and d.)

To investigate the importance of vascular systemic and local transport we took advantage of two existing *icals3m* lines driven from root phloem specific promoters: the *SISTER OF APPLE (SAPL)* promoter, which is specific to companion cells (plus meta sieve elements) and the *CALLOSE SYNTHASE 8 (CALS8)* promoter, which is expressed in phloem pole pericycle cells (Ross-Eliott et al., 2017). The cell-type expression pattern seemed conserved in inflorescence stems, based on sections from reporter lines. GUS signals delineated patterns resembling vascular strands (Fig. 1c) and *erYFP* signal was restricted to the phloem side of stem vascular bundles (Supp. Fig. 1c). In the case of *SAPL* the fluorescence signal was specifically associated with round cells smaller or of equal size to neighbouring sieve elements (the latter are identifiable by thick walls and associated strong calcofluor stain), while for *CALS8* fluorescence was observed in less regular and larger cells next to sieve elements (Fig. 1c; Supp. Fig. 1c). These patterns, when compared to electron micrographs of the inflorescence phloem (Nintemann et al., 2018) are compatible with *SAPL* expression in companion cells and *CALS8* in (a potential subset of) phloem parenchyma cells. The pericycle, as an anatomical structure, is absent in above ground tissues (Dubrovsky et al., 2001). However, as xylem pole pericycle like cells have also been described in above ground tissues (Sugimoto et al., 2010), it is plausible that an equivalent domain exists in stems, raising interesting questions about its function. Fluorescence-activated nucleus sorting and laser capture microdissection data in a transcriptome of the inflorescence stem of Arabidopsis (Shi et al., 2020) further supported *SAPL* and *CALS8* expression in phloem cell types and enrichment in the phloem cap, although in that case expression in other tissues (which we did not observe) was also reported (Supp. Fig. 1a, b). The promoters we use in the reporter constructs and in the *icals3m* constructs are the same, enabling correlations between the two.

Branching assays are performed using inflorescence stem explants, carrying one or more nodes, placed in Eppendorf tubes sealed with parafilm (Supplementary methods). Estradiol, required to induce the *icals3m* constructs, was basally supplied in the liquid medium in which the explants were placed. This approach had successfully induced *BRC1* expression in buds (Seale et al., 2017). To validate activation of the *icals3m* constructs we tested *CALS3* expression levels in the inflorescence stem following 24h of induction. Normalised expression roughly doubled in *SAPL:icals3m* and *CALS8:icals3m* but not in Col-0 (Supp. Fig. 3a). We then assessed whether this transcriptional induction resulted in callose accumulation in the stem, as expected. While some fluorescence signal from secondary antibodies raised against callose is to be expected, even in non-induced conditions, strong fluorescence signal was generally restricted to individually spaced cells (Fig. 1d; Supp. Fig. 3b). Strong signal is likely to correspond to sieve elements (Ross-Eliott et al., 2017). Clear callose signal was instead observed in clusters of neighbouring cells 48h after estradiol supply. The accumulation was specific to the phloem part of vascular bundles (Supp. Fig. 3b) and matched well the expected domains of expression of the promoters with *SAPL:icals3m* driving strong signal in smaller cells, and *CALS8:icals3m* driving callose deposition in larger cells (Fig. 1d).

In roots, induced callose accumulation in *CALS8:icals3m* (but not in *SAPL:icals3m*) results in blocked phloem unloading before and at 24h. This could be relevant in the context of bud growth as sucrose and Tre6P have been suggested to play roles in axillary bud activation (Fichtner et al., 2017; Mason et al., 2014; Fichtner et al., 2020). Axillary buds were collected at 0, 24 and 72h from explants bearing two nodes and with intact apices placed in eppendorf tubes. Cytokinin was supplied basally to the medium in which the explants were placed to enable bud escape from apical dominance and hence bud activation (Muller et al., 2015), likely via increased PIN3,4,7 levels in stem membranes (Waldie and Leyser, 2018) and downregulated *BRC1* levels (Braun et al., 2012; Dun et al., 2012; Seale et al., 2017). Variation in sucrose and Tre6P levels (normalised by protein content) could be observed between genotypes, treatments and buds of the same explants (bottom vs top buds). However, no consistent differences were observed between treatments. The magnitude of variation in the *icals3m* lines was similar to that of Col-0, which is not responsive to estradiol. The same trends were also observed in a repeat where top and bottom buds were pooled during collection (Supp. Fig. 4). These results could indicate that phloem unloading to buds is not affected in the *icals3m* lines employed (at time points when callose accumulation is visible in our sections) or that buds can modulate and buffer metabolite levels regardless of phloem delivery. Overall, these types of analyses can only provide static pictures of metabolites and don’t describe fluxes. In our hands it was not possible to perform phloem transport assays in inflorescence explants. Cell-cell communication might also be only transiently blocked by callose deposition as feedback mechanisms would eventually restore some level of trafficking. In either case, sucrose and Tre6P don’t seem severely affected upon callose accumulation so their levels might not underpin potential early bud growth differences in these treatments.

To study if the induction of callose had physiological effects on axillary bud growth we employed both inflorescence explants with intact apices carrying one or two axillary buds and explants with decapitated apices and two buds. In the latter case bud activation is intrinsically induced by reduced competition for auxin export (Crawford et al 2010). In all the experiments, buds from the *SAPL:icals3m* line reached shorter final mean lengths upon estradiol supply (Fig. 2). When two buds were present, the effect was generally more visible in the one growing more strongly (longer bud), irrespective of its position on the explant (top vs bottom bud). Whether the top/bottom bud or both activate in 2-node explants is not fully predictable, so we used a longer/shorter classification at the end of the time course. Traces for individual buds and plots showing relative growth biases between top and bottom buds are provided in Supp. Fig. 5. The lack of growth changes in Col-0 (and *CALS8:icals3m*) supports the notion that estradiol is not generally detrimental to bud growth (Fig. 2). The growth effect in *SAPL:icals3m* was not absolute or overwhelmingly strong and it was more pronounced in explants bearing two buds. It is easier for buds to grow in 1-node systems as competition occurs solely with the shoot apex while in 2-node explants three apices compete, making growth more sensitive to treatments.

**Fig. 2.**
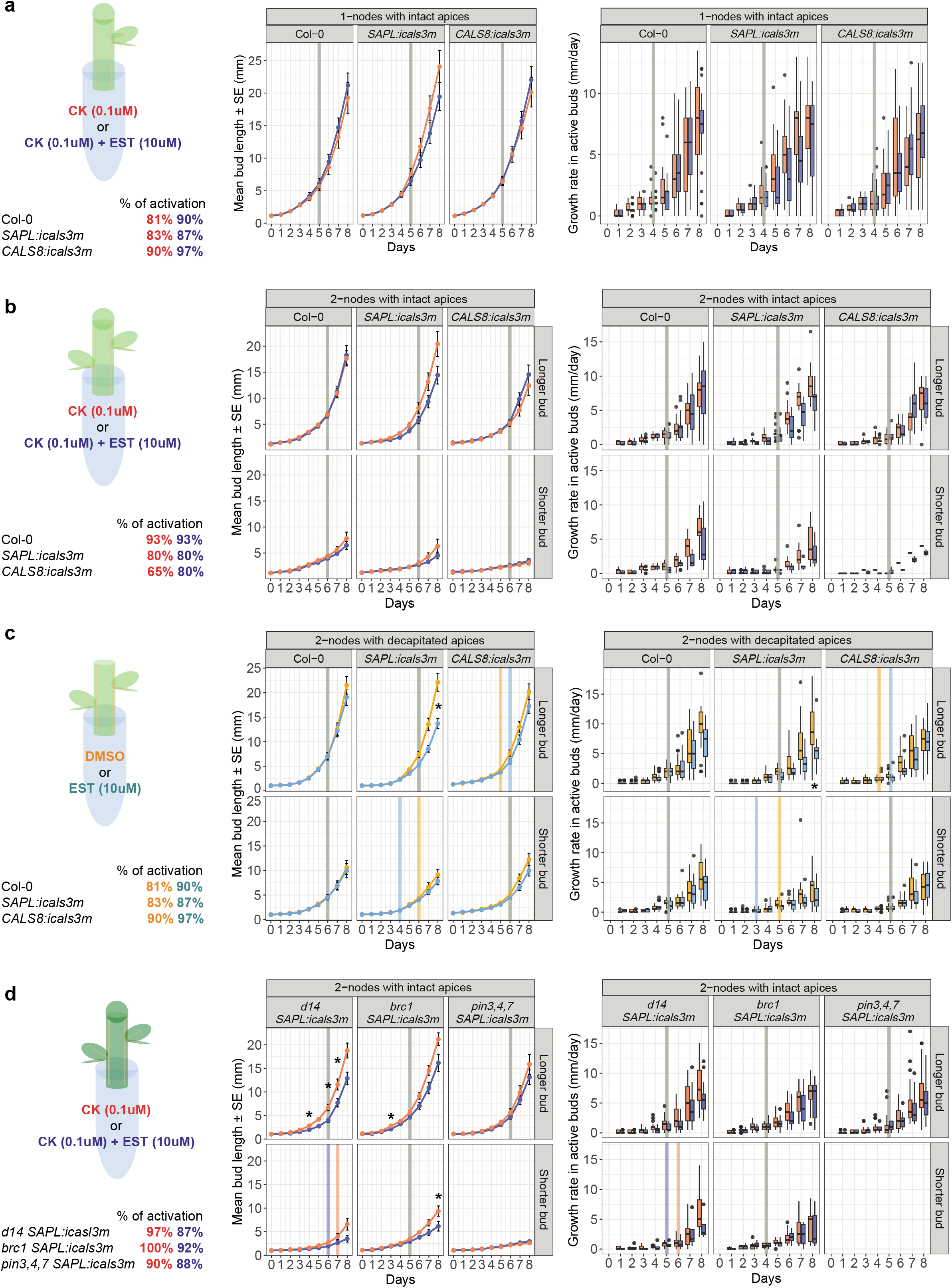
Activation percentages, mean bud lengths, median day of activation and growth rates upon mock/callose induction in the explant system employed. a. 1-node explants. b. 2-nodes explants with intact apices. c. 2-nodes explants with decapitated apices. d. 2-node explants from mutant genotypes with intact apices. Vertical lines indicate the median day of activation of the buds in each treatment (rounded to the nearest day). If the bars overlap they are coloured in grey, otherwise they follow the colour scheme employed throughout the figure. * indicates statistically significant differences in two-sided wilcox tests performed between treatments at each timepoint with a p-value threshold of <0.05. P-values, adjusted for multiple testing with the FDR method. N= 20-30 for all treatments and genotypes.

To identify what processes might be altered in the *SAPL:icals3m* line, three metrics were extracted to describe bud growth dynamics (Supplementary methods). First, the percentage of explants with at least one active bud, determined using a 5 mm length threshold at the end of the time course, was largely unaffected in all genotypes upon estradiol supply and was mostly above 75% (Fig. 2). Second, a proxy for the day of bud activation was calculated as the point of intersection of two linear regressions fitted to the growth curves of active buds (Fig. 2; Supp. Fig. 6). This metric was also largely unaltered upon estradiol treatment in the genotypes tested. The third metric, the growth rate of the active buds, seemed reduced at late timepoints, around and after the calculated day of activation (Fig. 2). This suggests that cell-cell communication within the *SAPL* domain might influence post-activation bud growth dynamics rather than activation itself.

The *SAPL:icals3m* line was then crossed with the Arabidopsis branching mutants *d14* (Waters et al, 2012), *brc1* (Gonzalez et al., 2013) and *pin3,pin4,pin7* (Bennett et al., 2016). We employed two node explants with intact apices to study if the construct’s effects were modulated in these genetic backgrounds. Despite being prone to bud activation and growth (Seale et al., 2017; Bennett et al., 2016) *d14* and *brc1* mutants still displayed the reduced elongation observed in a WT background. Growth was also generally restricted to the bottom bud in *d14* and *brc1* backgrounds (Supp. Fig. 5). Conversely, estradiol did not seem to elicit strong effects in the *pin3,pin4,pin7* background. PIN proteins, which are known to be involved in bud activation, might therefore also play roles in the less studied process of post-activation growth. Interplays between cell-cell communication and hormonal networks have already been reported in other contexts (Paterlini, 2020). It is important to point out that while our focus was targeted occlusion of PD, the observed extensive callose deposition in the wall of the *SAPL-*expressing cells (likely beyond the immediate PD positions)(Fig. 1) could also somehow impact apoplastic signals.

These results are also interesting when compared to the role of PD in seasonal bud dormancy in perennial plants (Rinne et al., 2001). Reductions in cell-cell permeability (via callose) are observed as buds enter endodormancy and these changes need to be relieved before buds can start to grow in spring (Singh et al., 2019; Tylewicz et al., 2018; Rinne et al., 2011). Phloem long distance transport is also blocked during dormancy via callose plugs (Aloni and Peterson, 1997). Cell-cell communication, therefore, seems central for bud (re)activation in endodormancy species. In paradormant plants like Arabidopsis, where growth inhibition is mainly imposed by signals associated with other organs on the plant rather than external clues (Lang et al., 1987), the role of PD seems to instead pertain to post activation growth rates.

Overall, the data shown here suggest that symplastic communication, involving companion cells of the inflorescence, might provide levels of regulation for the growth of a bud via unknown mechanisms. Indeed, the callose-related changes observed in *SAPL:icals3m* seem unrelated to early sugar levels in buds (preceding growth differences) and to the presence of the D14 protein, which were candidates based on phloem-mobile bud regulators (Mason et al., 2014; Fichtner et al., 2017; Kameoka et al., 2016). The *SAPL* domain might be the source or the receiver of an unknown regulator (potentially a protein or RNA) that might need to be loaded/unloaded in companion cells during long distance transport or be trafficked locally. The possibility that *SAPL* expressing cells in the shoot perform phloem unloading or enable metabolite fluxes can’t be entirely excluded from the bud metabolite levels we measured here. In addition to specific points raised before, pooling of buds that might have different growth dynamics could have partially influenced the measurements. Sustained callose production could also become a competing sink for local metabolites, which might be particularly relevant during late rapid bud growth. An unknown apoplastic unloading component could also eventually be influenced by callose accumulation. Despite these complication, since the *SAPL:icals3m* line rather than the *CALS8:icals3m* line displays phenotypic effects, which is the opposite of the situation in roots (Ross-Eliott et al., 2017), a reversal of importance of the respective cell types might be suggested in the shoot.

## Acknowledgements

We thank Jung-ok Heo and Sofia Otero (Sainsbury Laboratory - Cambridge) for providing *SAPL:erYFP* and *SUC2:erYFP* seeds respectively. We acknowledge Firas Bou Daher and Matthieu Bourdon (Sainsbury Laboratory - Cambridge) for their advice and help with immuno-localisations. We thank Emmanuelle Bayer (Laboratoire de Biogenèse Membranaire - Bordeaux) for critical reading of the manuscript. This work was supported by the Gatsby Foundation (GAT3395/PR3 grant awarded to YH and GAT3272C to OL) and the Max Planck Society (FF).

---

*Supp. Fig. 1. Phloem unloading into inflorescence stems and their organs*. a. SUC2:erYFP signal in sections across the vasculature in inflorescence stems. YFP is rendered in green, calcofluor white stain is false coloured in red and bright field is visible grey. b. *SUC2:GFP* signal in 7 days old roots c. and d. Zoomed panels from b. displaying the root tip and a differentiated part of the root. GFP is rendered in green, Propidium iodine stain is false coloured in red. Scale bars are of 50um and 10um (a), 500um (b), 200um(c-d). E. Absolute fluorescence or radioactive signal intensity in inflorescence organs of plants labelled with exogenous probes (N=17) relative to background signal in unlabelled plants (N=14). * Indicates statistically significant differences in non parametric, two-tailed Mann-Whitney tests with p-value thresholds of <0.05. f. Pooled fresh weight of the organs of inflorescences (N=17) of plants grown in jars in three repeats. Inferred weights in single inflorescences are calculated. g. Percentage of total fluorescent or radioactive signal (scaled by weight) in inflorescence organs. Letters indicate statistical differences in a Dunn’s test with a p-value threshold of <0.05.

*Supp. Fig. 2. Expression domains of SAPL and CALS8 promoters*. a. FANS data for the two promoters from Shi *et al*., 2020. b. LCM data for the two promoters from Shi *et al*., 2020. Intensities were scaled across each column to provide a picture of relative transcriptional enrichments of the promoters in the various tissues. These are not absolute levels of expression. c. *SAPL:erYFP* and *CALS8:erYFP* fluorescent signals in vascular bundles of the inflorescence stem. YFP is rendered in yellow while calcofluor white stain is false coloured in magenta. Zoomed panels specifically display phloem parts of vascular bundles. * indicates a SE cell. Scale bars are of 20um and 10um.

*Supp. Fig. 3. Induction of SAPL:icals3m and CALS8:icals3m constructs in inflorescence explants*. a. Expression levels of *CALS3* relative to control genes in inflorescence stems from explants treated for 24h with mock (DMSO) or EST solutions. * indicates a statistically significant difference in a two tailed Student’s t-test with a p-value threshold of <0.05. Callose immuno-labeling in inflorescence stem sections from explants supplied with DMSO or EST for 48h. Signal from secondary antibody is rendered in green while calcofluor white stain is false coloured in red. * indicates a cell with strong callose-related signal. Zoomed panels specifically display phloem parts of vascular bundles.

*Supp. Fig. 4. Metabolite levels in buds of 2-node explants with intact apices*. Sucrose and Trehalose-6P amounts at various time points after EST or non inductive treatment. In this repeat top and bottom buds from each explant were pooled together during collection. * indicates statistically significant differences in a two-tailed Mann-Whitney test with a p-value threshold of <0.05. N= 3-5 per timepoint, genotype and treatment.

*Supp. Fig. 5. Individual growth traces of Col-0, SAPL:icals3m and CALS8:icals3m buds upon EST or non inductive treatments and bottom-top bud relative growth biases in 2-node explants*. a. 1-node explants. b. c. 2-nodes explants with intact apices. d. e. 2-nodes explants with decapitated apices. f. g. 2-nodes mutant background explants with intact apices. Black solid lines represent mean lengths. In bias plots lengths at day 8 are marked with black dots. Diagonal of plot, indicating equal lengths, is shown as a dashed line. N= 20-30 for all treatments and genotypes.

*Supp. Fig. 6. Day of activation in Col-0, SAPL:icals3m and CALS8:icals3m inflorescence explants upon EST or non inductive treatment*. a. 1-node explants. b. 2-nodes explants with intact apices. c. 2-nodes explants with decapitated apices. d. 2-nodes mutant background explants with intact apices. Violin plots display distributions of data. Black horizontal bars indicate median values. Pairs of buds from the same 2-node are linked by dashed lines between the individual points. * indicates a statistically significant difference between treatments in KS tests with a p-value threshold of <0.05. P-values were adjusted for multiple testing with the FDR method. N= 20-30 for all treatments and genotypes.

