## Supplementary material for "Blocking plasmodesmata in specific phloem cell types reduces axillary bud growth in *Arabidopsis thaliana*": Materials and methods

**Supplementary methods**

*Fluorescent probe transport assays*

Plants grown in sterile jars were employed for experiments when their inflorescence bolts were around 15 mm in length. The plants were grown in a tissues culture room (16h light, 100 umoles m-2 s-1, 20°C, ambient humidity) in a 0.22% Murashige and Skoog Basal Salts, 0.05% 2-(N-morpholino)ethanesulfonic acid (MES), 1% sucrose, 1% Agar, pH 5.7 medium. To one leaf of each rosette 10 ul of a 10% Adigor (Syngenta) solution in water were applied. Adigor is an oil based adjuvant that facilitates the uptake of substances by leaves (Knoblauch et al., 2015; Knox et al., 2018). After two hours, the adjuvant was removed and immediately replaced with 10 ul of a 1 mM Carboxytetraethylrhodamine (CTER)(Acros Organics) solution in water and acetonitrile (Knoblauch et al., 2015). The probe was allowed to translocate within the plant for approximately 15h (overnight). Each inflorescence was then dissected and the different components (buds, shoot apex, stems and cauline leaves) placed in individual eppendorf tubes containing 450 ul of 100% ethanol. Tubes were labelled in order to identify components derived from the same inflorescence. The tubes were incubated for 1h at 75°C in a heating block. Samples were then briefly cooled on ice and spun down in a centrifuge at maximum speed for one minute. Two 200 ul aliquots from each eppendorf, without any plant material, were deposited in neighboring wells in a clear 96 well microplate (Greiner Bio-one). The aliquots acted as technical repeats. Plates were read in a SpectraMax i3x scanner (Molecular devices). To scale signals by fresh mass organs from equivalent inflorescences, grown in jars, were collected and weighted.

*Sucrose transport assays*

The plant material and the Adigor treatment were the same as described fluorescent probe transport assay. In place of the CTER probe, 10 ul of a C14-sucrose (Perkin Elmer) diluted in water was applied to the treated leaf (Activity in the solution: 0.00185 MBq/ul). The probe translocation, organ dissection and incubation in ethanol steps were the same as described above. The only difference was that 400 ul of 100% ethanol, rather than 450 ul were employed. The contents of each eppendorf, including the plant material itself, was transferred to 4 ml plastic vials (Perkin Elmer) already containing 600 ul of -40 Microscint scintillation liquid (Perkin Elmer). The vials were briefly vortexed at low speed to remove bubbles and mix the content. The samples were placed in the holder, inserted into the MicroBeta2 microplate counter machine (Perkin Elmer) and radioactive decay signals measured from the bottom of the vials for 2 min. Non-labelled controls were also included in the experiment.

*Heatmaps*

The tissue expression heatmap was generated with the online tool Heatmapper (Babicki et al., 2016).

*One and two node branching assays*

Arabidopsis plants were grown in plastic pots (one plant per pot) until the reproductive bolting stem was approximately 1-2 cm tall. The plants were grown in a Conviron growth chamber under long day conditions (16h light, 250 umoles m-2 s-1, 21°C during the day, 17°C in the night, 65% humidity). Lidless microcentrifuge tubes (1.5 ml) were filled with Arabidopsis thaliana solution (ATS) without sucrose (Wilson et al., 1990), supplemented as appropriate, and covered with a parafilm lid with a 1-2 mm hole in it. For intact assays stem segments bearing the primary shoot apex and two axillary buds (2-nodes) were excised and directly placed into the tubes. For decapitation assays the main shoot apex was removed above the uppermost of the two buds, using a needle under a Zeiss Stemi 2000 dissecting scope, before placing the stem segments into the tubes. The solutions for intact assays were ATS without sucrose, supplemented with 0.1 uM 6-benzylaminopurine (BA) and 10 mM beta-estradiol (EST), or supplemented with 0.1 uM BA and an amount of DMSO equal to that of EST. Concentrated stock solutions of EST and BA were dissolved in DMSO. The solutions for decapitation assays were ATS without sucrose supplemented with 10 mM EST or an equivalent amount of DMSO. For assays involving stem segments bearing the primary shoot apex and only one axillary bud (1-nodes), the full inflorescence was initially placed in tubes containing ATS solution alone. The stems were left to elongate for 3 days, separating the nodes sufficiently to allow the stem to be trimmed away below the most apical cauline node. The resulting explants were moved to new tubes with the same solutions as those used for 2-node assays with intact apices. In all cases stem segments were only selected when the axillary buds were shorter than 2 mm. The tubes with the stem segments were placed in Conviron growth chambers inside closed boxes to maintain humidity. Branch lengths were measured daily, to the nearest 0.5 mm, using a 30 cm metal ruler. Measuring started as soon as stem explants were placed in tubes (with the exception of the 1-node assays). The liquid in the tubes was topped up daily with a syringe. Dried out explants were discarded and their data excluded from analyses.

For the data analysis three metrics were extracted from the bud dynamics. First, the percentage of explants with active buds (or with at least one active bud if more were present) determined using bud lengths threshold of 5 mm at the end of the time course. Buds above this threshold were considered active. Second, the breakpoint day (a proxy for the day of activation of the buds) corresponded to the point of intersection of two linear regressions fitted to the bud growth curves. It was assumed that the curve shape can be simplified to two consecutive linear segments with different slopes. The linear segments were fitted using the *segmented* package (Muggeo, 2008) in R (R Core Team, 2017). A difference in length is required for breakpoint estimation and the metric was only calculated for (active) buds above 5 mm in length at the end of the time course. Third, the daily growth rate of active buds (above 5 mm at the end of time course) was also calculated as millimeters growth per day.

The code supporting these analyses can be found at the following GitHub repository: https://github.com/mvanrongen/publication_2020_Paterlini_plasmodesmata

*RNA extraction and cDNA synthesis*

Around 10 stems were collected from explants in the 2-node setup and pooled for each sample time point or treatment. When a zero hour time point was included, the buds were directly collected from the soil grown plants. They were harvested at a developmental stage equivalent to when explants would have been placed in 2 node set ups. Three to four biological replicates were collected for each timepoint or treatment and stored at -80°C. A metal or glass bead was added to each tube and samples were ground in a TissueLyser II machine (Qiagen). The RNA from these powdered samples was immediately extracted using the RNeasy Plant Mini Kit (Qiagen), following product specifications. Beta-mercaptoethanol and RLT buffer were employed in the protocol. The extracted RNA was then treated with the TURBO DNA-free Kit (ThermoFisher Scientific), following product specifications, in order to remove any contaminant DNA. Lastly, to generate the cDNA, 1 ug of RNA was added to 1 ul of Oligo(dT)12-18 primers (ThermoFisher Scientific), 1 ul of dNTP Mix (10 mM each) and distilled water up to 12 ul total volume. The mixtures were heated to 65°C for 5 min and quickly chilled on ice before adding 4 ul of First-strand Buffer, 2 ul of Dithiothreitol (0.1M) and 1 ul of RNaseOUT Recombinant Ribonuclease Inhibitor (ThermoFisher Scientific). The mixtures were then incubated at 42°C for 2 min and then 1 ul of SuperScript II Reverse Transcriptase (ThermoFisher Scientific) added. Lastly, the mixtures were incubated for 42°C for 50 min and then the reactions were inactivated by heating at 70°C for 15 min. The obtained DNA was stored at -20°C.

*qPCR*

The cDNA samples were diluted 20x in water. A 384 multiwell plastic plate (Roche) was prepared by hand. In each well 0.8ul of 10uM forward primer, 0.8ul of 10uM reverse primer, 4.4ul of water, 10ul of SensiFAST No-Rox Sybrgreen (Bioline) and 4ul of diluted cDNA were added (total volume 20ul). Thermocycling was carried out using a Lightcycler 480 (Roche). Sequences of the primers are listed in this table.

| **Gene** | **Primer** | **Sequence** | **Reference** |
|---|---|---|---|
| *CALS3* | FW | GTTGGGGGATGCTCTTGA | Vaten *et al.*, 2011 |
| *CALS3* | RV | ACGAGCTAGGGTCCTCACTG | Vaten *et al.*, 2011 |
| *SANDF* (Control) | FW | CCAAGATACAACGCTCAGGC | Czechowski *et al.*, 2005 |
| *SANDF* (Control) | RV | CAAGGCGTACCCTGCAATCT | Czechowski *et al.*, 2005 |
| *PP2A* (Control) | FW | GCGGTTGTGGAGAACATGATACG | Czechowski *et al.*, 2005 |
| *PP2A* (Control) | RV | GAACCAAACACAATTCGTTGCTG | Czechowski *et al.*, 2005 |

The Cycle threshold (Ct) values were calculated using the Second Derivative Maximum analysis function included in the Lightcycler 480 software. Melt curves were also determined for all reactions carried out. Technical repeats were discarded if melt curves clearly contained more than one peak or if the Ct value for that repeat was more than 0.5 units away from the other repeats with the same mix (pipetting issues). Ct values were then converted into an expression value, using the formula 2-Ct, and averaged across technical replicates. Relative expression levels were then calculated comparing the expression of the gene of interest to the geometric mean of the expression of the control genes being used. The values were displayed as log2.

*Sugar content extraction*

Around 20 buds were collected from explants in the two-node setup and pooled for each sample time point or treatment. Three or four biological replicates were collected. Samples were flash frozen and stored at -80°C until further processing. The samples were held in a plastic rack, in a basin filled with liquid nitrogen and 175 ul ice-cold chloroform:methanol (3:7 v/v) were added to each eppendorf tube. The liquid freezes. The tissues were ground with a pestle as the mixture gradually thawed. The pestle was rinsed with 175 ul of ice cold chloroform:methanol (3:7 v/v), which were then added to the corresponding sample. Samples were left at -20°C for 2h with occasional mixing. To extract the sugars, 350 ul of ice cold water was added to each sample and the samples were allowed to warm up to 4°C by repeated shaking. Samples were centrifuged at 13000 g, at 4°C for 10 min. The upper aqueous phase was transferred to a new tube and kept on ice. The lower phase was re-extracted adding 300 ul of water, mixing and centrifuging. The second upper aqueous phase so obtained was combined to the first one and these samples (containing the sugars) were evaporated to dryness in a centrifugal vacuum (miVac Quattro concentrator - GeneVac) at 35°C. The lower phase in the original tubes (containing, among other substances, proteins) were also evaporated to dryness. The analysis protocols followed those described in Fichtner et al. (2017); Lunn et al. (2006). Sugar abundances were determined by liquid chromatography–mass spectrometry while protein content was determined colorimetrically using dye-binding assays.

*Fixation protocol*

Bolting stem segments from 2-node setups or equivalent inflorescences taken directly from soil-grown plants were collected. They were snap fixed in eppendorf tubes in a solution of 2.5% formaldehyde, 2.5% glutaraldehyde in Phosphate-buffered saline (PBS) 1x within a Pelco Biowave Pro Microwave Tissue Processor (Ted Pella Inc) for 10 min at 175 watts, under vacuum conditions, at 26°C. Samples destined for immuno-labelling were then additionally left in the fixative overnight at 4°C, all other samples were immediately processed for sectioning.

*Vibratome sectioning*

Fixed material was washed twice in PBS and then placed in individual wells of a plastic casting tray. Warm 2% agarose was pipetted over the samples, to fill the wells. Tweezers were used to gently arrange the samples in vertical or horizontal positions depending on the sectioning plane of interest. Once solidified, the agarose sample cubes were gently removed from the casting tray and superglued to the sample holder of a Leica VTI200S vibratome. Once the samples had firmly attached to the holder, water was added to fill the sample container of the machine, submerging the block. The block was then sequentially cut with a double edged razor blade (Wilkinson) oscillating at an amplitude of 1 mm and at a speed of 0.5 mm/s. The thickness of the sections was set at 200 um. Sections were gently fished out of the water and placed in shallow, water filled, containers for further processing or imaging.

*GUS staining*

To assess GUS activity in the sections, they were placed in an X-Gluc solution (30 mM disodium phosphate, 20 mM monosodium phosphate, 0.5mg/ml X-Glu, 0.1% v/v Triton X-100, 0.5 mM potassium ferrocyanide, 0.5 mM potassium ferricyanide in water) and incubated in darkness at 37°C until blue colouring became visible on the tissue. Incubation time varied depending on the promoter of interest. The staining solution was then removed and replaced with multiple washes of 70% ethanol (with incubation at 37°C) until the tissue had cleared. Images were then acquired using the SPOT5.2 imaging software and a Leica M165 FC Stereo Microscope, in its bright field mode, with an attached SPOT pursuit USB camera.

*Immuno-labelling*

Sections from different biological samples were returned to separate wells in the plastic casting trays. Three biological samples per treatment were used. To each well 1 ml of 2% Bovine serum albumin (BSA) in PBS was added. The samples were then placed in the Pelco Biowave Pro Tissue Processor and treated for 1 min at 170 watts, under vacuum conditions, at a temperature of 26°C. Samples were then left under vacuum for 20 min. The BSA solution was replaced in each casting well with 600 ul of primary antibody solution: 0.25 ul / 100 ul of a monoclonal mouse antibody against (1-3)-beta-D-glucan (callose) (Biosupplies Australia) in PBS. Samples were treated for 1 min in the Pelco Biowave Pro Microwave Tissue Processor with the same settings listed above. Samples were then left in the vacuum for 2h. The primary antibody solution was replaced by 1 ml of 2% BSA in PBS. Four additional BSA washes were performed using the Pelco Biowave: 1 min each with the setting listed above. The BSA solution was ultimately replaced with 600 ul of the secondary antibody solution: 0.25 ul / 100 ul of F(ab')2-Goat anti-Mouse IgG (H+L) Cross-Adsorbed Secondary Antibody, Alexa Fluor 488 (Thermo-Fisher Scientific). Samples were treated for 1 min with the same settings listed above. Samples were then left in the vacuum for 2h. As before, the antibody solution was replaced by 1 ml of 2% BSA in PBS and four additional BSA washes were performed using the Pelco Biowave Pro Microwave Tissue Processor: 1 min each with the setting listed above. Sections were left in PBS overnight and the next day mounted in a solution of Calcofluor-white M2R (Sigma-Aldrich) and Citifluor anti-fading solution 1 (Citifluor): 10 ul of 100x calcofluor in 495 ul water and 495 ul anti-fading solution.

*Confocal microscopy*

For immuno-labelled sections a Zeiss LSM700 microscope was used. Sections were excited with a 488 nm laser wavelength for GFP and 405 nm for Calcofluor white counterstain. For all other confocal imaging purposes a Leica SP8 microscope was employed. A 405 nm and a 514nm laser were used to excite Calcofluor white and YFP respectively. Images were acquired with 25x or 63x water dipping objectives. The GFP laser intensity and corresponding channel gains were kept constant during each experiment and the signal was never saturated. Images had a 1024x1024 pixels format and were acquired using line averaging of 5. Maximum intensity projections of Z-stacks were generated in the Fiji imaging software (Schindelin et al., 2012).
