## Supplementary material for "Blocking plasmodesmata in specific phloem cell types reduces axillary bud growth in *Arabidopsis thaliana*": Supp. Fig. 1

**a**

*SUC2:erYFP*

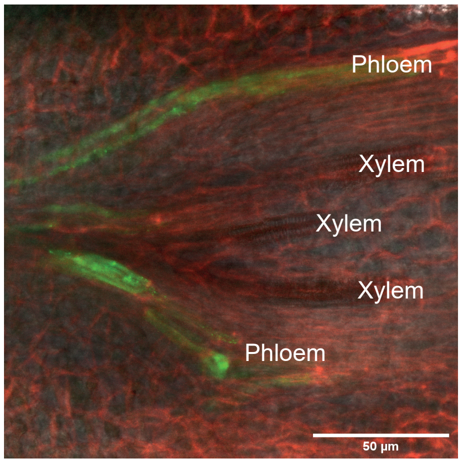


**b**

*SUC2:GFP*

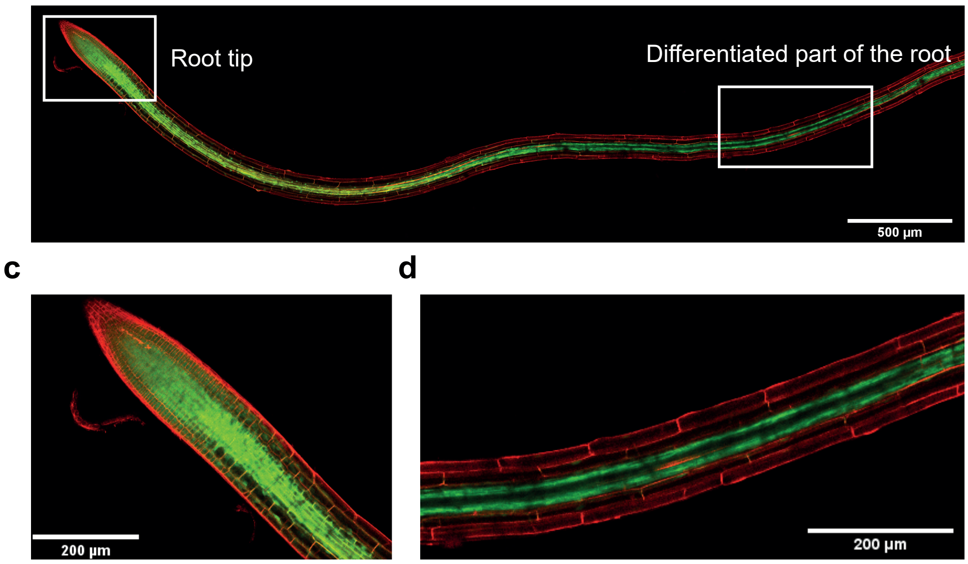


**e**

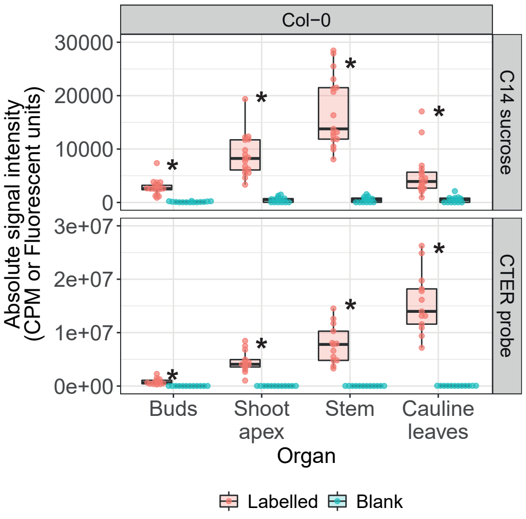


**f**

| Rep | Organ | Pooled weight | Individual weight |
|---|---|---|---|
| 1 | Buds | 4. 6 mg | 0.3 mg |
| 1 | Shoot apex | 8.1 mg | 0.5 mg |
| 1 | Stem | 11.5 mg | 0.7 mg |
| 1 | Cauline leaves | 20.4 mg | 1.2 mg |
| 2 | Buds | 5.5 mg | 0.3 mg |
| 2 | Shoot apex | 7.9 mg | 0.5 mg |
| 2 | Stem | 9.3 mg | 0.5 mg |
| 2 | Cauline leaves | 25.3 mg | 1.6 mg |
| 3 | Buds | 3 mg | 0.2 mg |
| 3 | Shoot apex | 8.1 mg | 0.5 mg |
| 3 | Stem | 7.6 mg | 0.4 mg |
| 3 | Cauline leaves | 27.6 mg | 1.6 mg |

**g**

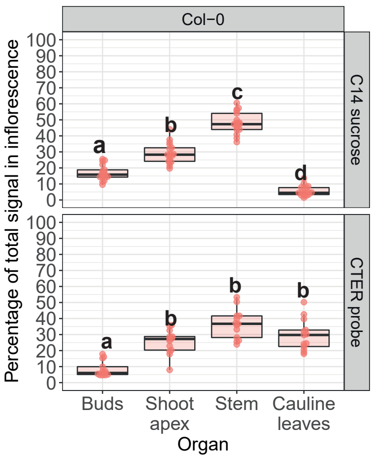


**Supp. Fig. 1**
