## Supplementary figures and images for "Blocking plasmodesmata in specific phloem cell types reduces axillary bud growth in *Arabidopsis thaliana*"

### Supp. Fig. 2

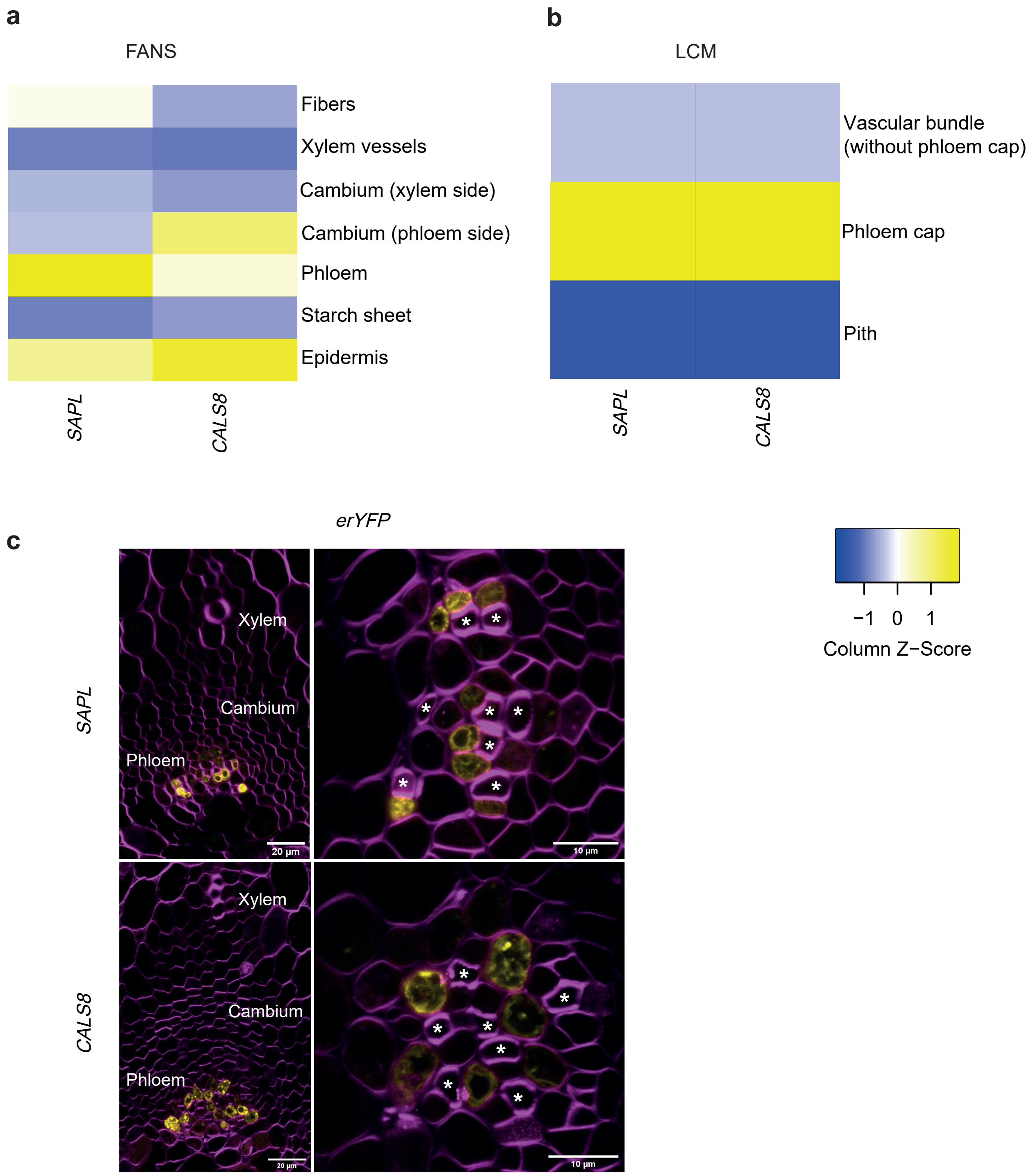

### Supp. Fig. 3

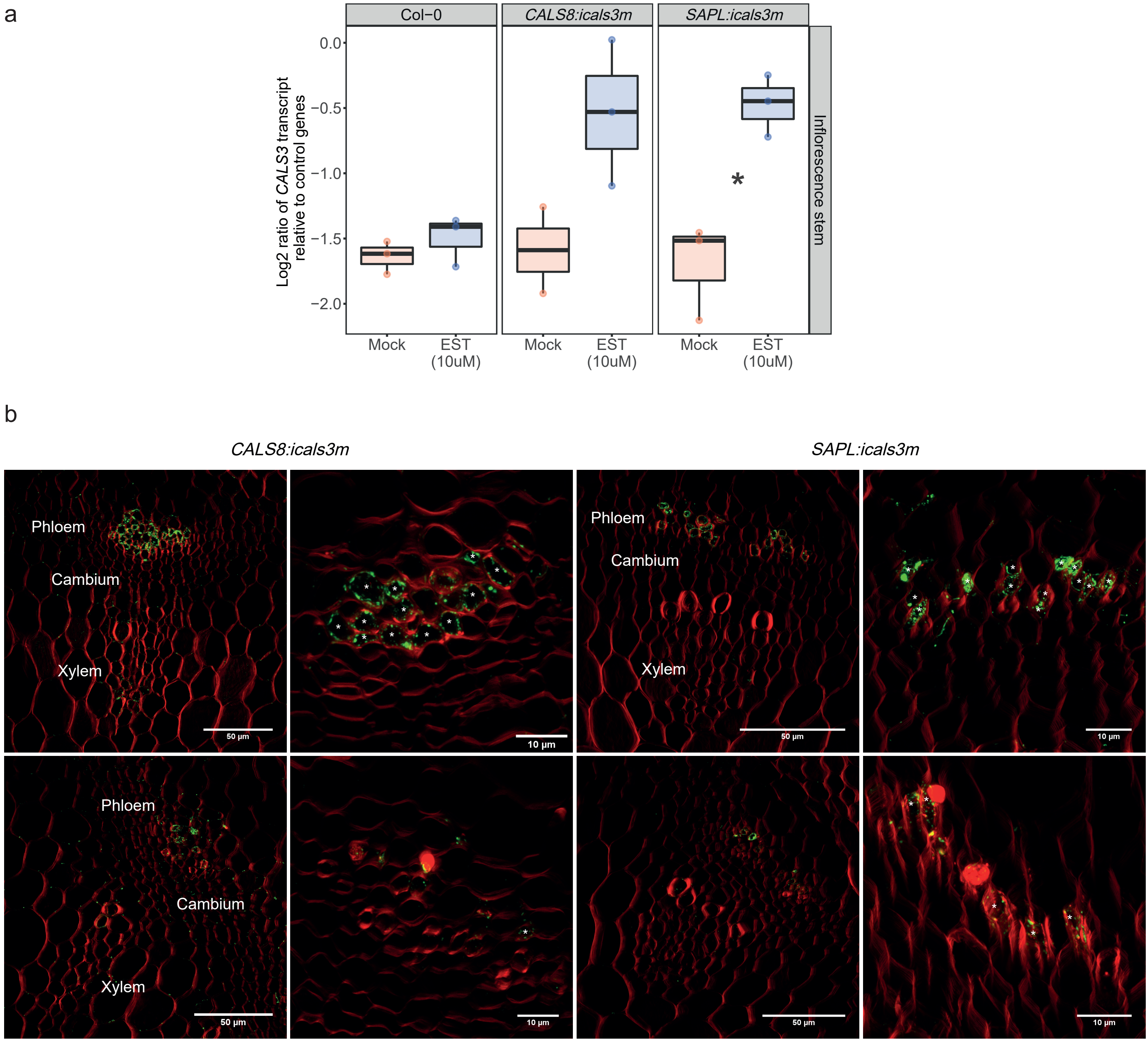

### Supp. Fig. 4

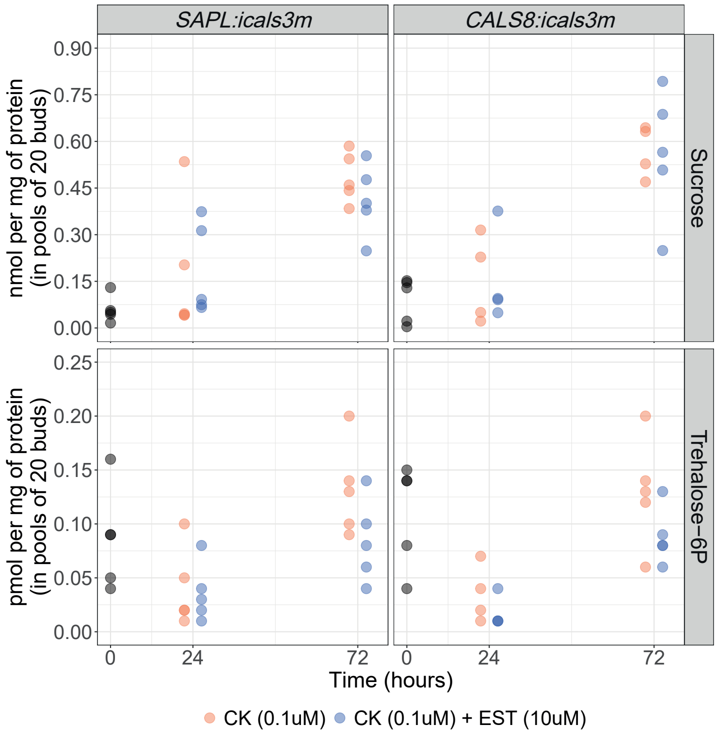

### Supp. Fig. 5

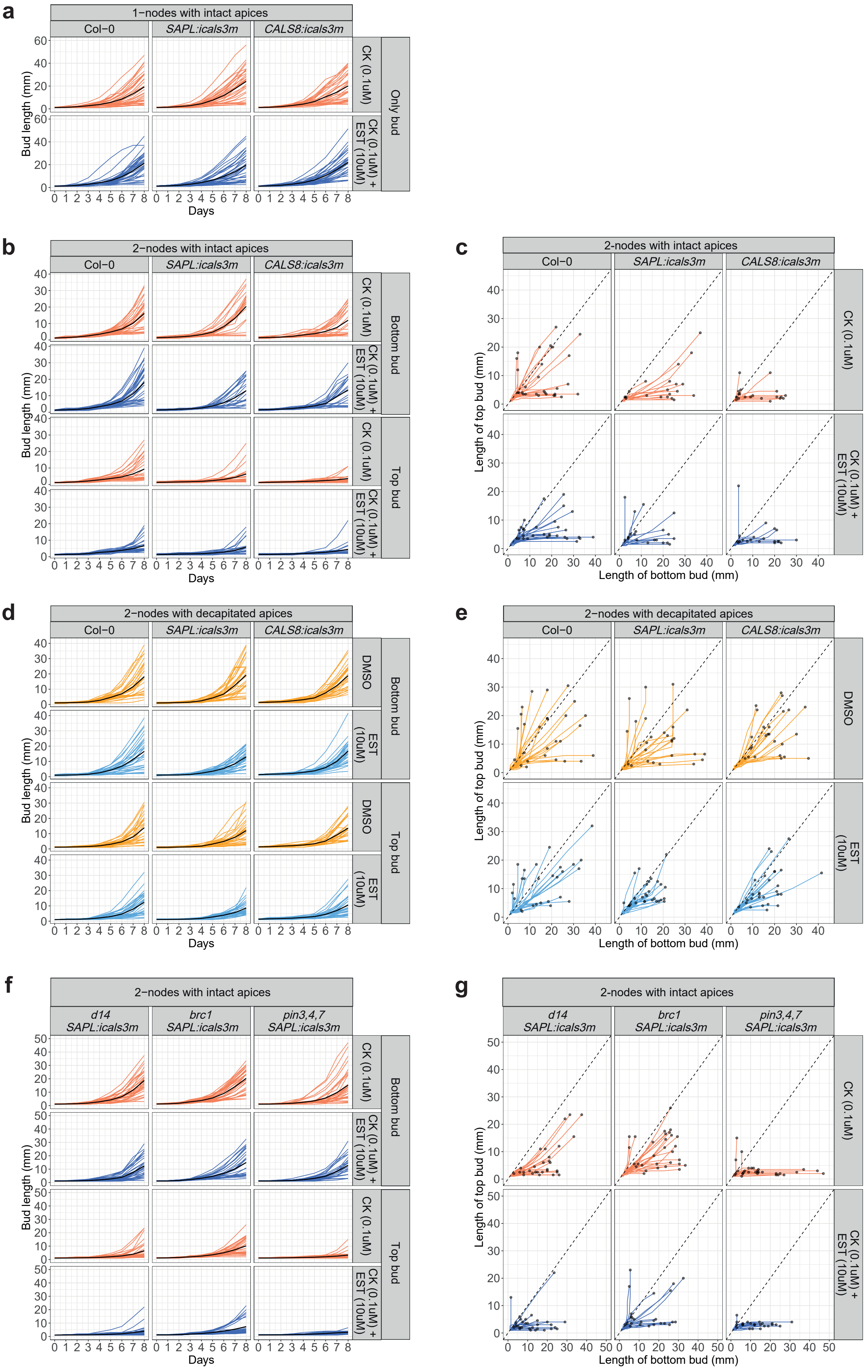

### Supp. Fig. 6

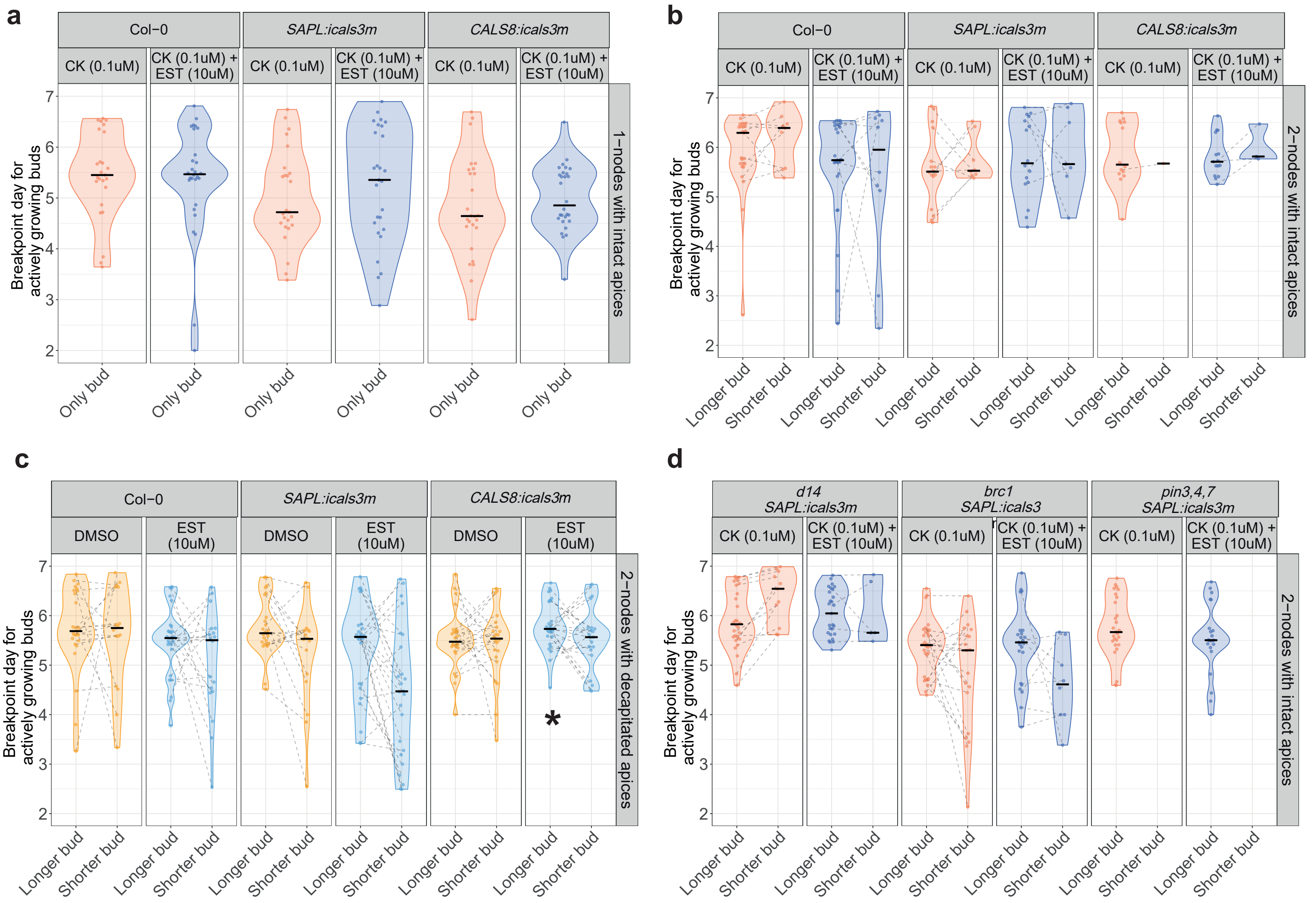
